# Revealing the microbial heritage of traditional Brazilian cheeses through metagenomics

**DOI:** 10.1101/2022.03.14.484326

**Authors:** Caroline Isabel Kothe, Nacer Mohellibi, Pierre Renault

**Author notes:** Corresponding authors: C. I. Kothe. INRAE, Micalis Institute, Domaine de Vilvert, 78350 Jouy-en-Josas., P. Renault. INRAE, Micalis Institute, Domaine de Vilvert, 78350 Jouy-en-Josas.

## Abstract

Brazilian artisanal cheeses date from the first Portuguese settlers and evolved via local factors, resulting in unique products that are now part of the patrimony and identity of different Brazilian regions. In this study, we combined several culture-independent approaches, including 16S/ITS metagenetics, assembly- and deep profiling-metagenomics to characterize the originality of the microbiota of five varieties of Brazilian artisanal cheeses from the South and Southeast regions of Brazil. Their core microbiota contained mainly lactic acid bacteria (LAB), of which *Lactococcus lactis* subsp. *lactis* was the most frequent, followed by *Streptococcus thermophilus* in the South region. Moreover, several samples from the Southeast region contained, as dominant LAB, two other food Streptococci belonging to a new species of the *salivarius* group and *S. infantarius*. Rinds of samples from the Southeast region were dominated by the halotolerant bacterium *Corynebacterium variabile* and the yeasts *Diutina catenulata* and, to a lesser extent, by *Debaryomyces hansenii* and *Kodamaea ohmeri*. Rinds from the South region contained mainly LAB due to their short ripening time, and the predominant yeast was *D. hansenii*. Phylogenomic analysis based on *L. lactis* metagenome-assembled genomes (MAGs) showed that most Brazilian strains are closely related and form a different clade from those whose genomes are available at this time, indicating that they belong to a specific group. Lastly, functional analysis showed that *S. infantarius* acquired a ∼26 kb DNA fragment from *S. thermophilus* starter strains that carry the LacSZ system, allowing fast lactose assimilation, an adaptation advantage for growth in milk. Finally, our study identified several areas of concern, such as the presence of somatic cell DNA and high levels of antibiotic resistance genes in several cheese microbiota, implying that the milk used was from diseased herds. Overall, the data from this study highlight the potential value of the traditional and artisanal cheese production network in Brazil, and provide a metagenomic-based scheme to help manage this resource safely.

## 1 Introduction

Beyond their historical, socio-economic and cultural aspects, artisanal fermented foods have recently received increased attention as potential sources of biological resources that can provide both technological and health benefits. Among them, traditional cheeses worldwide contain rich natural microbiota that improve the quality of cheese production in terms of flavor, texture and safety aspects (Montel et al., 2014; Walsh, Macori, Kilcawley, & Cotter, 2020). This is also true for emerging countries where dairies have more recently appeared.

There are few reports on the colonization of cheese in Brazil. Some studies have suggested that the potential origin of Brazilian dairy products traces back to the “discovery” of the country in 1581 when Portuguese settlers arrived with their cattle and passed on cheese production techniques to the local population (Borelli, Lacerda, Penido, & Rosa, 2016; Penna, Gigante, & Todorov, 2021). Although cheese production in Brazil was based on European practices, factors such as climate, farm organization, feed, breed of dairy animals, quality of raw milk and indigenous microbiota resulted in unique products that are now part of the patrimony and identity of different Brazilian regions (Kamimura, Magnani, et al., 2019; Penna et al., 2021). There are more than 30 varieties of artisanal cheeses produced in micro-regions of the Southeast (Araxá, Campo das Vertentes, Cerrado, Canastra and Serro), South (Colonial and Serrano), Center (Caipira), North (Marajó) and Northeast (Butter and Curd) regions of the country (Kamimura, Magnani, et al., 2019). Recent studies have shown that these local cheeses are potential sources of lactic acid bacteria (LAB) that can be used for the development of starter cultures (Cabral et al., 2016; Margalho et al., 2020). It could thus be interesting to more widely explore the biodiversity of these artisanal products whose microbiota may have evolved separately from the well-studied Western cheeses. These data would be valuable to more effectively develop them as biological resources, ensure their safety and facilitate their preservation, as well as to support the development of technical regulations and maintain their historical and cultural footprint.

Several studies have focused on the microbiological characterization of Brazilian artisanal cheeses using culture-dependent methods (Lima, Lima, Cerqueira, Ferreira, & Rosa, 2009; Luiz et al., 2016; Perin, Sardaro, Nero, Neviani, & Gatti, 2017; Pontarolo et al., 2017; Resende et al., 2011). However, these techniques do not provide a complete picture of the existing microbiota since several species are poorly cultivable or require appropriate culture media. Recently, Kamimura *et al*. (2019) (Kamimura, De Filippis, Sant’Ana, & Ercolini, 2019) conducted a pioneering study using culture-independent methods in Brazilian cheeses. Using 16S rRNA sequencing, they characterized the bacterial diversity of a wide number of artisanal cheeses from different regions of Brazil at the genus level. They mainly identified LAB of the genera *Enterococcus, Lactobacillus, Lactococcus, Leuconostoc* and *Streptococcus*, as well as some probable contaminants such as *Enterobacteriaceae* and *Staphylococcus*. However, these studies seem to focus on the core microbiota, leaving the rind microbiota unknown, whereas it may harbor halotolerant and halophilic bacteria and yeasts, which are key actors in cheese ecology and technology (Banjara, Suhr, & Hallen-Adams, 2015; E. Dugat-Bony et al., 2016; Kothe et al., 2021; Wolfe, Button, Santarelli, & Dutton, 2014). Finally, the 16S gene analysis does not provide functional data or detailed taxonomic information that make it possible to uncover the original characteristics of their microbiota.

Consequently, there are still many unanswered questions concerning these artisanal Brazilian cheeses that would require deeper investigation, including species-and even strain-level analysis, for yeasts as well, and that target the specific microbiota of the rind and the core. While metagenetics allows a rapid identification at the genus level, other meta-omic approaches such as shotgun metagenomics may provide a more accurate microbiota level of analysis. In this study, we focused our analyses on cheeses from two Brazilian regions (South and Southeast) and combined 16S rRNA and ITS metagenetic analysis with shotgun metagenomic analysis in order to reveal their specificities at the level of species composition and strain origin, and to highlight functional features of interest. These data could be a benefit to market expansion, ensuring the protection of historical and sanitary aspects, in addition to supporting the development of technical regulations, legislation and standardization parameters for the different types of traditional Brazilian cheeses.

## 2 Methodology

### 2.1 Sample collection and DNA extraction

A total of 23 artisanal cheeses were obtained from Southeast and South Brazil from five varieties: Araxá, Canastra, Serro, Colonial and Serrano. Samples were collected from local producers, artisan markets and fairs, and sent by post to France. The rinds were separated from the cores using sterile knives, and both fractions were analyzed to obtain a more detailed overview of the microbial diversity of those cheeses. The samples were diluted 1:1 (w/v) in guanidinium thiocyanate 4M solution (Sigma-Aldrich, USA) with Tris-HCl 0.1M, and mixed in an Ultra Turrax T25 (Labortechnik) at 8,000 rpm for 2 min. A 10% N-lauryl sarcosine solution (Sigma-Aldrich, USA) was added to the mixture, vortexed and centrifuged at 4°C and 14,000 rpm for 30 min. The fat and supernatant were eliminated and the pellet was used for DNA extraction using the protocol described by Almeida *et al*. (2014) (Almeida et al., 2014). DNA quality was visualized on 0.8% agarose gel and the quantification was measured with a Qubit 2.0 fluorometer (Life Technologies) using a Qubit dsDNA HS (High Sensitivity) Assay Kit.

### 2.2 Metagenetic analyses

Bacterial diversity was analyzed by sequencing the amplified region V3-V4 of the 16S rRNA gene using primers V3F (5’-ACGGRAGCWGCAGT-3’) and V4R (5’- TACCAGGGTATCTAATCCT-3’). Additionally, fungal diversity was evaluated in rinds using ITS3 (5’-GCATCGATGAAGAACGCAGC-3’) and ITS4 (5’-TCCTCCGCTTWTGWTWTGC-3’) primers. The PCR was performed with MTP Taq DNA Polymerase (Sigma-Aldrich, USA), and the cycling conditions were: 94°C for 1 min, followed by 30 cycles of amplification at 94°C for 1 min, 65°C for 1 min, and 72°C for 1 min, with a final extension step of 10 min at 72°C. The sequencing was performed with a V3 Illumina MiSeq kit, as described in Poirier *et al*. (2018) (Poirier et al., 2018).

The quality of the raw data was evaluated with FastQC (Wingett & Andrews, 2018) and the sequences were imported into the FROGS pipeline (Escudié et al., 2018) to obtain the Operational Taxonomic Units (OTUs). The sequences were filtered by length (150–500 bp) and then pooled into OTUs with SWARM (Mahe, Rognes, Quince, de Vargas, & Dunthorn, 2014) with the distance parameter of 3. Chimeras were removed with VSEARCH (Rognes, Flouri, Nichols, Quince, & Mahe, 2016) and OTUs with at least 0.01% in the whole dataset were retained. The OTUs were affiliated with SILVA 132 SSU databases (Quast et al., 2013) for bacteria and UNITE 8.2 for fungi (https://unite.ut.ee/). Alpha-diversity and beta-diversity analyses were performed in R Studio v.3.6.1 using the phyloseq and ggplot2 packages (v1.30.0) (McMurdie & Holmes, 2013; Poirier et al., 2018).

### 2.3 Taxonomic composition by shotgun metagenomics

The DNA of 15 cheese samples was sequenced using Illumina HiSeq2500 technology at GATC-Biotech (Konstanz, Germany), which yielded between six and eight million paired-end reads of 150-nucleotide length. Metagenomic reads corresponding to the *Bos taurus* genome were filtered with Bowtie2 (Langmead & Salzberg, 2012) and visualized with Samtools flagstat (H. Li et al., 2009). From the generated fastq files, we first estimated microbial composition by mapping the samples reads against the representative clade-specific marker catalogue contained in the MetaPhlAn tool, v.3.0.4 (Truong et al., 2015).

Additionally, we performed taxonomic profiling using an assembly-based marker gene analysis, which allows non-supervised binning of metagenomes. To do this, the reads were trimmed for quality and *de novo* assembly was performed using metaSPAdes, v.3.9 (Bankevich et al., 2012). Genes were then predicted using Prodigal (v.2.6.3) and marker genes were extracted using fetchMG, v.1.0 (Ciccarelli et al., 2006; Sunagawa et al., 2013). We chose to perform our taxonomic assignations by using the *ychF* marker gene, whose closest homologue was assigned by a blastp search on all the available sequences from the NCBI protein database. Summary species composition plots were created in R (v.3.6.1) using the ggplot2 package, v.3.3.2. Finally, phylogenetic analyses were performed with the *ychF* proteins identified using ClustalX 2.1 (Thompson, Gibson, Plewniak, Jeanmougin, & Higgins, 1997) and MEGA7 (Kumar, Stecher, & Tamura, 2016) with the Neighbor-Joining method (Saitou & Nei, 1987) and 1,000 bootstrap replicates (Felsenstein, 1985).

### 2.4 Metagenome-assembled genome (MAG) analyses

Genome binning was performed using MetaBAT2-2.12.1 (Kang, Froula, Egan, & Wang, 2015), with a minimum contig size of 1,500 nucleotides and the default settings. The quality of the resulting prokaryotic bins was assessed with CheckM (Parks, Imelfort, Skennerton, Hugenholtz, & Tyson, 2015), and MAGs < 80% completeness and/or > 10% contamination were excluded. For eukaryotes, we considered those bins that aligned with > 60% of their sequence length to fungal reference genomes from GenBank as MAGs of quality.

In addition, MAG assemblies were performed for Streptococcal species by additional binning, such as blastn analysis with reference genomes of related species. Contigs were then filtered by percentage of the length covered, identity and coverage levels. Finally, manual curing of questionable contigs (new, low coverage of homology on references, etc.) were performed by blastn and blastx analysis against nr/nt and nr NCBI databases, respectively.

### 2.5 Phylogenomic and functional analyses

The Automatic Multi-Locus Species Tree web server (https://automlst.ziemertlab.com/) (Alanjary, Steinke, & Ziemert, 2019) was used to determine closely related genomes based on core gene alignments of the recovered MAGs. The closest species were inferred based on the percentage of average nucleotide identity (ANI) calculated using FastANI, v.1.31 (Jain, Rodriguez-R, Phillippy, Konstantinidis, & Aluru, 2018).

kSNP (v.3.0) was used to perform the phylogenomic analysis for the *Lactococcus lactis* species (Gardner, Slezak, & Hall, 2015), with the maximum likelihood option and kmer size of 21. A total of 225 genomes of the *Lactococcus lactis* species were downloaded from Genbank and combined with our MAG to compute this analysis.

The search for antibiotic resistance (ABR) genes was performed by read mapping against the CARD database (Jia et al., 2017; McArthur et al., 2013) using the PATRIC web server (Antonopoulos et al., 2019). The presence of particular genes in reference genomes, such as virulence factors, ABR and genes of technological interest, was determined using the FoodMicrobiome tool (https://migale.jouy.inra.fr/foodMicrobiome/; (Kothe et al., 2021). This tool performs read mapping on given reference genomes and provides read counts for each annotated gene. Additionally, this tool was used to detect, with high sensitivity and reliability, the subdominant populations of given species by analyzing the distribution of metagenomic reads on reference genomes of these species.

### 2.6 Data availability

Raw sequences of amplification of 16S rRNA and ITS genes and raw metagenomic reads were deposited on the European Nucleotide Archive (ENA) under the BioProject ID PRJNA693797. The MAGs are available in: https://data.mendeley.com/datasets/4w8w9mkfjy/1

## 3 Results

### 3.1 Sample descriptions

A total of 23 artisanal cheeses were obtained from the Southeast and South regions of Brazil from five varieties: Araxá, Canastra, Serro, Colonial and Serrano (**Fig. 1**). The cheeses were produced with raw milk, except C-19, and the samples were collected from local producers, artisan markets and fairs (**Table S1**). These five cheese varieties are defined as semi-fat to fat (25-59.9% of fat) and semi-hard cheeses (humidity 36-45.9%) (de Medeiros Carvalho, de Fariña, Strongin, Ferreira, & Lindner, 2019; MAPA, 2020; RS, 2016). The differences in the production process and characteristics of each type of cheese are briefly described in **Supplementary Data 1**.

**Fig. 1.**
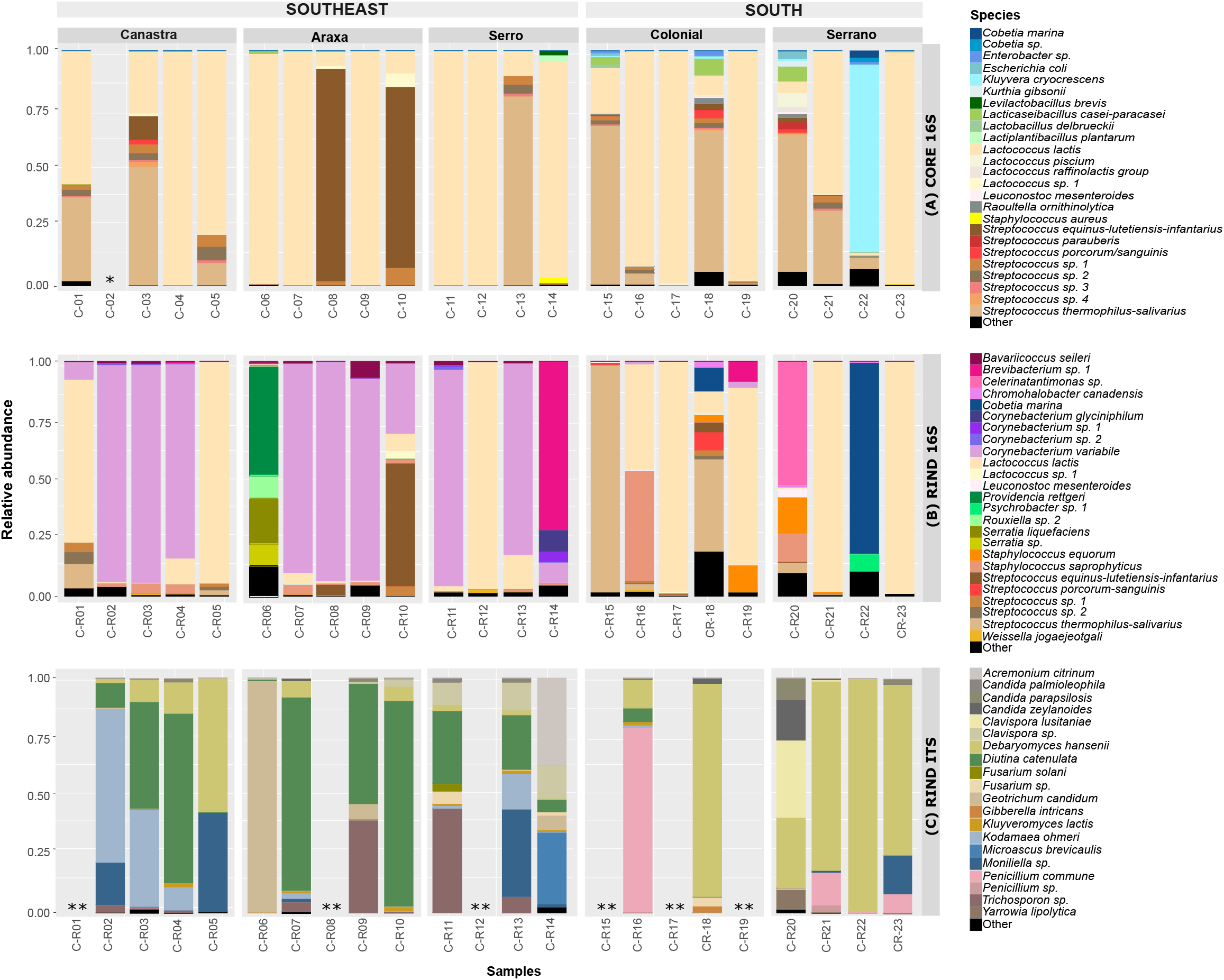
Map of Brazil showing regions and types of cheeses collected.

### 3.2 Taxonomic diversity of Brazilian cheeses using amplicon sequencing approaches

The bacterial diversity present in the core and the rind of the 23 Brazilian cheeses was assessed using amplicon sequencing targeting the 16S rRNA gene. The sequences were clustered in 100 bacterial OTUs whose taxonomic assignment was possible up to the species level in the majority of the cases (**Table S2**). The C-02 sample was discarded due to its low read depth. For the majority of other samples, the rarefaction curves reached the saturation plateau, indicating that the sequencing depth was sufficiently recovered (**Fig. S1)**.

The alpha-diversity analysis for core samples showed greater observed richness of bacterial species (p < 0.01) in the South samples compared to the Southeast samples (**Fig. 2A)**. However, the bacterial evenness measured by Shannon and inverse Simpson indices showed that the diversity of species in both regions was quite similar (p > 0.05). The Principal Coordinates Analysis (PCoA) and clustering showed the diversity among the samples, and we observed that the majority of cheese cores from the two regions shared a common bacterial microbiota (**Fig. 2A**). Nevertheless, the samples C-08 and C-10 from the Araxá micro-region contain microbiota that is different from the other samples but similar between them, and C-22 belongs to a particular group. Indeed, the taxonomic composition showed that LAB constitute the major part of core sample microbiota with *Lactococcus lactis* and *Streptococcus thermophilus*-*salivarius* as the dominant species (over 50% of the reads) in 14 and five samples, respectively, while a *Streptococcus* from the *equinus-lutetiensis-infantarius* complex is dominant in the C-08 and C-10 samples (**Fig. 3A**). Additionally, in the core of the C-22 sample, we observed the predominance (∼79.5%) of the enterobacteria *Kluyvera cryocrescens*, while *S. thermophilus* was present at only low levels (4.6%). Finally, several other LAB species were less abundant, such as *Levilactobacillus brevis, Lacticaseibacillus casei, Lb. delbrueckii, Lactiplantibacillus plantarum, L. piscium, Lc. mesenteroides*, as well as undefined species of *Streptococcus* and *Lactococus* (**Fig. 3A**).

**Fig. 2.**
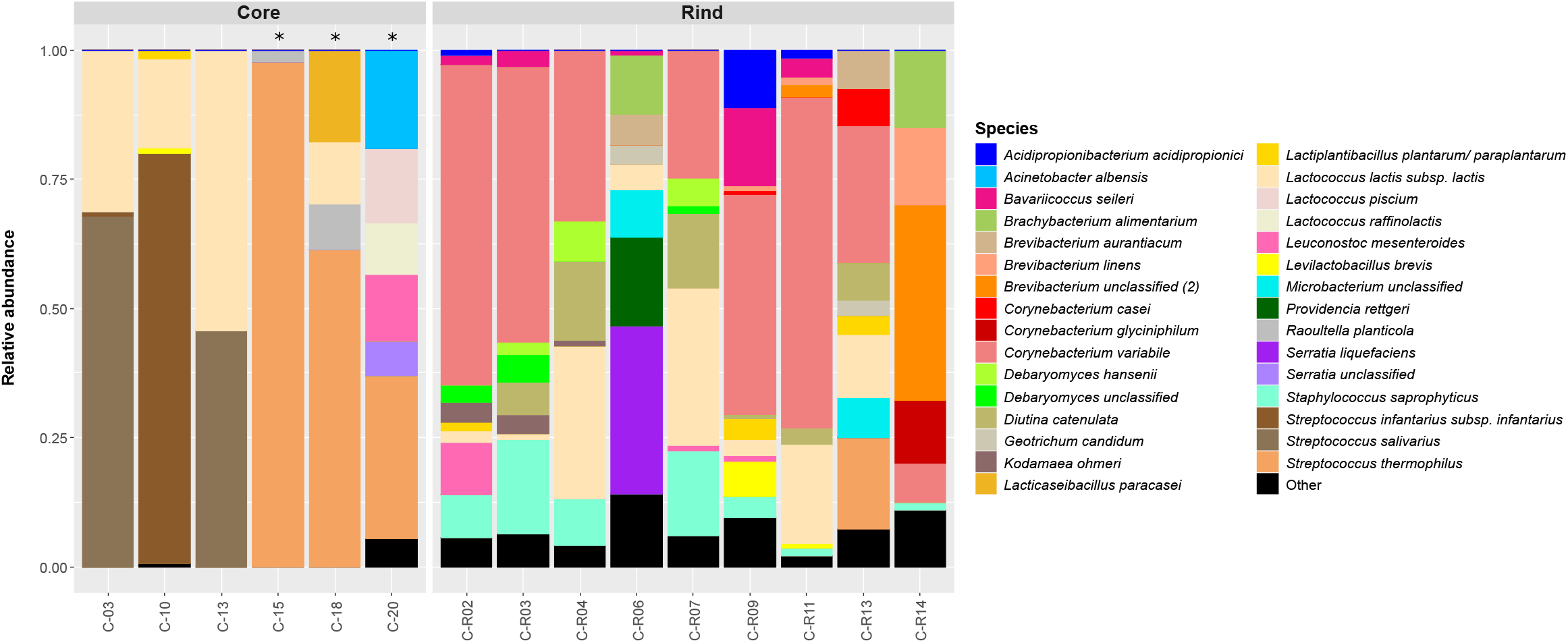
Boxplots of alpha-diversity indices, PCoAs and clustering among bacterial communities identified in the core **(A)** or rind **(B)** of Brazilian cheeses, and fungal communities identified in the rind **(C)**. Samples are colored according to the different regions where they are produced, i.e., blue for South and red for Southeast.

**Fig. 3.**
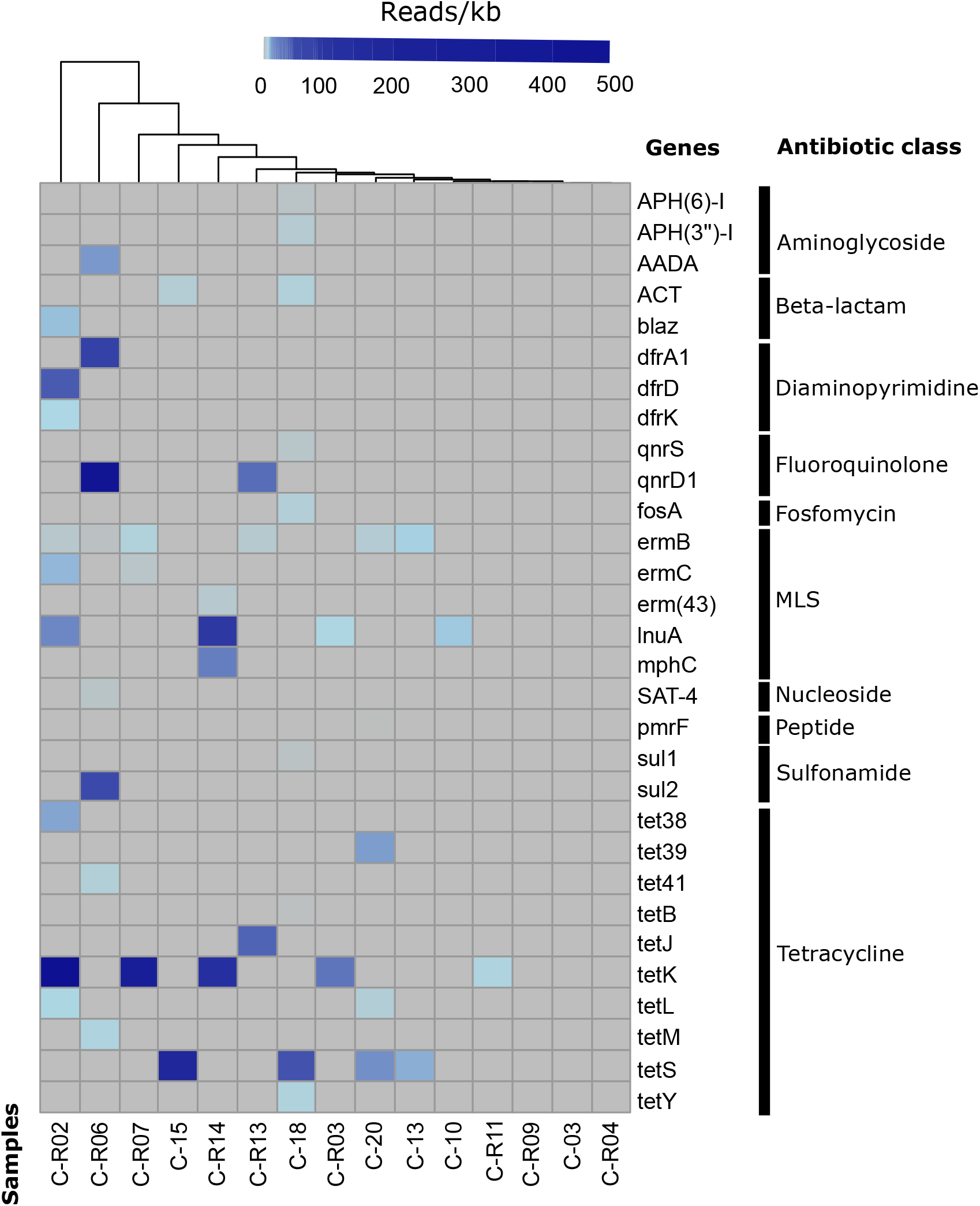
Bacterial (16S, **A-B**) and fungal (ITS, **C**) plot depicting the relative abundance of the 25 main species found in Brazilian cheese cores and rinds. One asterisk (*) indicates low depth of the sequences and two asterisks (**) indicate samples that are not amplified.

Concerning bacterial richness and evenness of the rind samples, no statistical differences were observed based on the observed species and Shannon and Simpson diversity indices (p > 0.05, **Fig. 2B**). The PCoA and hierarchical clustering mainly showed two different groups of microbiota: one composed only of samples from the Southeast and another with cheeses from both regions. The taxonomic composition showed that the majority of cheese rinds from the South displayed the same dominant species as their core (seven out of nine), whereas those from the Southeast region are dominated by *Corynebacterium variabile* (eight out of 14) (**Fig. 3B**). The two divergent samples from the South, C-R20 and C-R22, were dominated by high levels of *Celerinatantimonas sp*. and *Cobetia marina*, respectively. Finally, four of the divergent rind samples from the Southeast had a composition closer to that of the core (C-R01, C-R05, C-R10 and C-R12), and the last two, C-R06 and C-R14, were dominated by *Enterobacteriaceae* species and an undefined *Brevibacterium* species, respectively (**Fig. 3B**).

Furthermore, *S. aureus* and *E. coli* species, which may have pathogenic strains, were detected in more than 45% of core samples and more than 35% of rinds at levels > 0.01%. The highest levels were detected in C-14 (which represented 2.86% of *S. aureus*) and C-20 samples (with 3.45% of *E. coli*) (**Fig. 3A**). Concerning eukaryotic diversity, ITS amplification was successful for 17 rind samples, and the sequences were grouped into 34 yeast/fungal OTUs **(Table S2**). The alpha diversity analyses did not show statistical differences between the South and Southeast regions (p > 0.05). The beta-diversity analysis showed two groups, the first one consisting mainly of samples from the South, except C-R05, and the second one of samples from the Southeast (SE), except C-R16 (**Fig. 2C**). Taxonomical analysis of the ITS indicated the predominance of different fungal/yeast species between South and SE cheeses (**Fig. 3C**). Using a threshold of > 5% of read abundance, *Debaryomyces hansenii* was present in all South cheese rinds (n=6), but only in five from SE cheeses (45%). Conversely, the dominant yeast in SE cheeses was *Diutina catenulata*, which was present in nine samples at levels > 5%, and in only one South cheese. Moreover, several additional species were detected mainly in SE cheeses, such as *Kodamaea ohmeri, G. candidum, Trichosporon sp*. and *Moniliella sp*., and in South cheeses, such as *Penicillium commune, Clavispora lusitaniae* and *Candida zeynaloides*.

### 3.3 Refined microbiota by metagenomics

In order to more precisely analyze the microbiota from our samples, we selected six and nine samples of cores and rinds, respectively, to perform shotgun metagenomic analyses. Such analyses allow a taxonomic identification up to strain level and the simultaneous assessment of the level of reads corresponding to bacteria and fungi. Moreover, it makes it possible to detect eventual virulence factors in pathogenic species, as well as antibiotic resistance genes and genes of technological interest.

#### 3.3.1 Species level analyses

Metagenomic samples often contain animal reads, which may significantly bias the analysis when they are in great abundance. In the present samples, we detected *Bos taurus* reads at levels ranging from 0.1 to 76% (**Table S3**). We observed that cow reads were notably higher in the core samples, especially C-15, C-18 and C-20 from the South region, where they accounted for more than 30%.

The taxonomical composition of the 15 cheese metagenomes was first determined using MetaPhlAn, which yielded 128 bacterial and one fungal species (**Table S4**). Species composition obtained by this analysis is relatively in accordance with those obtained with 16S rRNA amplicon for species present at abundance levels > 1%. Interestingly, it resolved several taxonomical assignations such as that of *Streptococcus equinus-lutetiensis-infantarius*, which becomes *S. infantarius* and appears to be the dominant LAB in two Araxá cheeses (**Fig. 3A**). It also detected several species present at significant levels such as *Corynebacterium variabile, Staphylococcus saprophyticus, Brachybacterium alimentarium, Acidipropionibacterium acidipropionici, Brevibacterium linens* and *S. parauberis*. Conversely, it did not detect two possibly abundant bacteria, an uncharacterized *Brevibacterium* sp. and *Providencia rettgeri* in C-R14 and C-R06, respectively. Moreover, it detected only one yeast (*Candida parapsilosis*) in our metagenome set, whereas ITS amplicon analysis showed large fungal diversity, indicating a probable lack of corresponding genome references in the database currently available from MetaPhlAn.

In order to characterize microbiota independently of fixed references, we applied a marker gene analysis from the assembled metagenomes, which yielded 57 bacterial and seven fungal species (**Table S4**); the overall composition is shown in **Fig. 4**. Although the results are consistent with the previous analysis made by read profiling, some differences merit highlighting. In particular, 14 species whose relative abundance ranges from 3 to 37% were detected uniquely by the marker analysis, probably due to their absence in the currently implemented MetaPhlAn database. Among bacteria of potential technological interest, it revealed *Lacticaseibacillus paracasei* and two potential new species of *Brevibacterium* and one of *Microbacterium*. Moreover, this analysis suggested that while the *S. thermophilus*-*salivarius* populations detected in the South samples belong to *S. thermophilus*, those detected in the Southeast samples are instead related to another species of the *salivarius* group (**Table S4, Fig. S2**). Indeed, their sequence of the well-conserved *ychF* gene displays more than 5% nucleotidic divergence with those of the other species of the *S. salivarius* group, suggesting that they belong to a new species. Finally, we detected seven species of yeasts, *Debaryomyces hansenii, Diutina catenulata, Kodamaea ohmeri, Geotrichum candidum, Kluyveromyces lactis* and two other Saccharomycetales. We also identified the proportions of prokaryotes and eukaryotes in the samples; yeasts appear to be absent in the core samples, and range from 0 to 27% in the rinds.

**Fig. 4.**
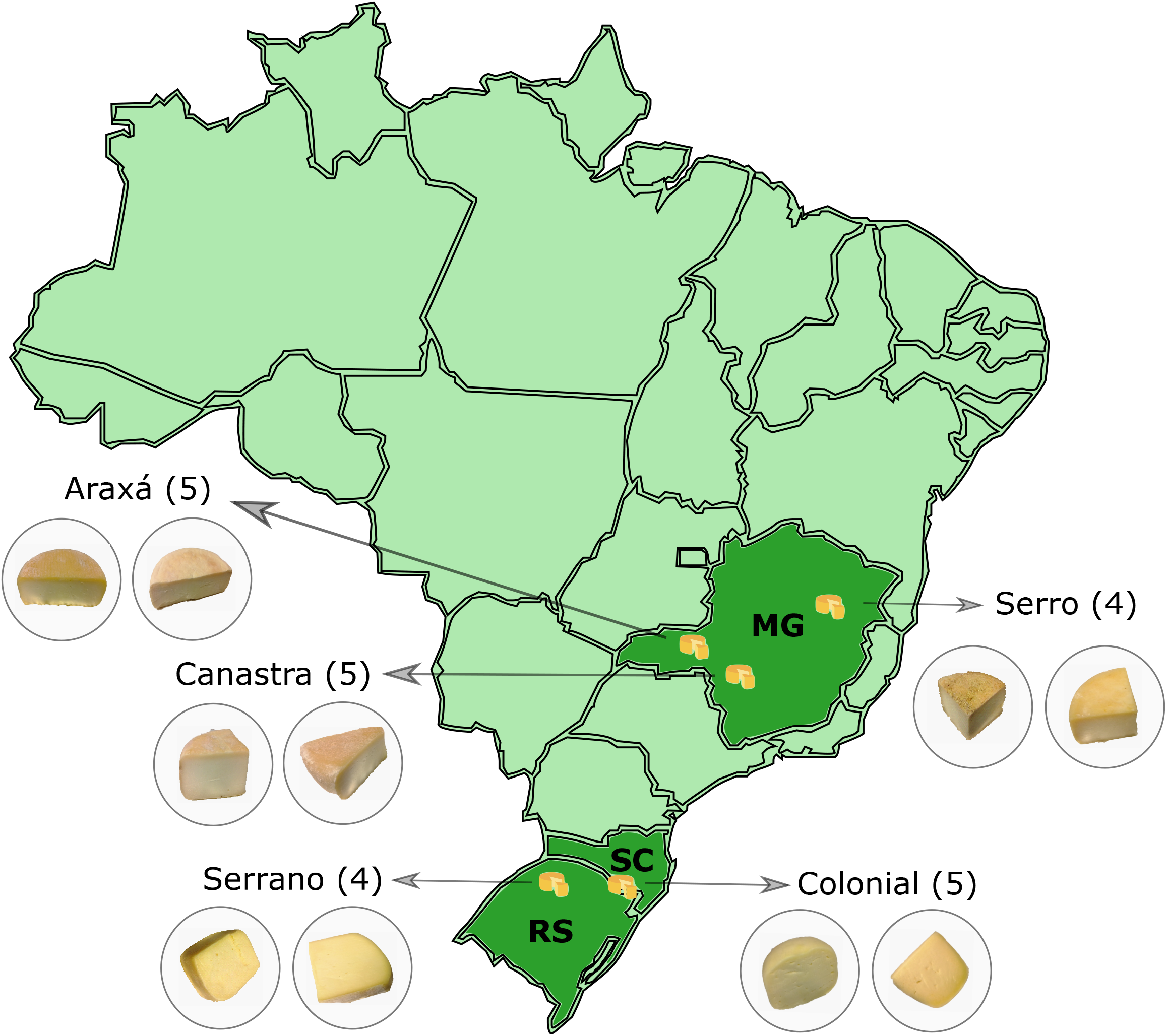
The relative abundance of microbial composition of 15 Brazilian cheeses (six cores and nine rinds). Values were calculated from the coverage of the *ychF* marker gene assembled in the metagenomes. The asterisk (*) indicates a low depth of the sequences.

#### 3.3.2 MAGs and strain-level analyses

A total of 54 prokaryotic and 10 eukaryotic metagenome-assembled genomes (MAGs) of high quality were reconstituted from the metagenome assemblies (**Table S5**). They include 27 prokaryotic and four eukaryotic species. The majority of bacterial MAGs correspond to *L. lactis* subsp. *lactis* (10) and *C. variabile* (8). Moreover, eight MAGs represent six potential novel species since they have less than 95% average nucleotide identity (ANI) with reference genomes (**Table S5, Fig. S3**). They also include species of the genera *Brevibacterium* (2), *Streptococcus, Corynebacterium, Lactobacillus* and *Microbacterium*. Additionally, one MAG that belongs to the *Micrococcaceae* family could not be identified at the genus level since it does not share ANI > 80% with any reference genome.

Further detailed analysis was performed with the Lactococcal and Streptococcal MAGs, which correspond to key species of starter bacteria. Phylogenomic analysis of the *L. lactis* MAGs was performed with 215 reference genomes of this species, representing strains from its two main subspecies (*lactis* and *cremoris*) and isolated from different environments (dairy, vegetable, animal, etc.) (**Fig. S4**). This showed that nine of the Brazilian cheese MAGs form a homogeneous group whose most closely related strains are *L. lactis* subsp. *lactis* isolated from various traditional dairy products around the world. Although several strains of this group are referenced at NCBI as belonging to the diacetylactis biovariant, they all lack the citrate lyase complex (*mae-citRCDEFXG*) involved in the production of diacetyl from citrate, and they are phylogenetically distant from the diacetylactis group. Additionally, the *L. lactis* MAG originating from the C-R02 sample is clustered with another *L. lactis* subsp. *lactis* group containing mostly industrial dairy starter strains. A search for the genes encoding the Lac-PTS system (ten genes) and PrtP that allow rapid lactose assimilation and casein breakdown in Lactococci, respectively, were found in all metagenomes at levels similar to or slightly higher than chromosomal Lactococcal genes (**Table S6**).

Concerning *Streptococcus*, six MAGs were built, and ANI analyses showed that they may belong to three species (**Table S5**). A first group of three MAGs belong to *S. thermophilus* species (ANI > 98% with the genome of reference strains of this species), and detailed phylogenomic analysis clearly links them to strains isolated from cheese and milk in Europe. A second group of two MAGs (C-03 and C-13) belongs to the *salivarius* group, forming a cluster clearly separated from *S. thermophilus* and related to but different from *S. salivarius* (ANI∼94%, **Fig. S3A**). This result indicates that they probably belong to a new species. Finally, MAG-C-10, belonging to the species *S. infantarius*, was compared to the genomes of the type strain isolated from feces and the CJ18 food strain. The three genomes share over 98.5% ANI and 1,576 orthologous proteins, confirming their close relationship. Moreover, MAG-C-10 displays a contig of 26.1 kb, sharing 99.6% identity over 25,870 nucleotides with *S. thermophilus* ATCC 19258. This region contains, in particular, the *lacZ* and *lacS* genes that allow the rapid assimilation of lactose of *S. thermophilus*. It is flanked at the 5’ end by genes belonging to the IS3 mobile element and at its 3’ end by a 55-nucleotide signature flanking the IS1182 family transposase in Streptococci. Complementary analysis by read mapping showed that this region is covered at the same average level as the 1,516 genes defined above as being common to the three *S. infantarius* strains.

Concerning species growing on the rinds, eight MAGs referring to *C. variabile* were recovered. They display a very close relatedness (ANI > 99%) with the only two genomes deposited at the NCBI (DSM 44702 and Mu292 strains isolated from smear-ripened cheeses in Ireland (Schröder, Maus, Trost, & Tauch, 2011) and France (E. Dugat-Bony et al., 2016), respectively. Furthermore, MAGs corresponding to known cheese halotolerant bacteria such as *S. saprophyticus* (three MAGs), *Bavaricoccus seileri* (4), *Brachybacterium alimentarium* (2) and several *Brevibacterium* species were recovered.

The ten fungal MAG reconstituted in this study could be assigned to four species: *Debaryomyces hansenii, Diutina catenulata, Geotrichum candidum* and *Kodamaeae ohmeri* (**Table S5**). Genomic comparison with the reference strains available in the NCBI database shows that *G. candidum* and *K. ohmeri* MAGs display ANI > 99% and alignment to reference > 80% to CLIB918 and 148 strains isolated from Pont l’Evêque cheese in Normandy (France) and from honeybee gut in the USA, respectively. Finally, *D. catenulata* and *D. hansenii* MAGs present ANI = ∼97.5% and alignment to reference of 60-90% (depending of the MAG quality) and 95%, respectively.

#### 3.3.3 Detection of pathogens, virulence factors and antibiotic resistances

Strains of 12 pathogen species commonly found in dairy products were searched for by deep profiling analysis in the Brazilian cheese metagenomes (**Table S7**). *S. aureus* was detected at significant levels in the Canastra cheese rind sample C-R02 (0.5% of reads mapped), with an average gene detection rate of 14.6 reads/kb. Since this level of coverage allows a reliable detection of genes in metagenomes, we searched for the presence of 27 staphylococcal enterotoxins responsible for foodborne outbreaks, and only enterotoxin A (*sea, entA*) and X (*selx*) were found. Finally, among the other species, only *Escherichia coli* was detected in two core samples from the South (C-15 and C-20), at average levels of 1.3 and 2.5 reads/kb, which is below the threshold required to perform a reliable detection of virulence genes.

In addition, a mapping-based approach against a comprehensive collection of antibiotic resistance genes was performed. A total of 30 ABR genes were identified, belonging to nine different classes of antibiotics (**Fig. 5** and **Table S6**). The presence of ABR genes was detected in all samples, except in three (C-R09, C-03 and C-R04). The C-R02 cheese sample displayed the highest number (nine ABR genes). Overall, the most abundantly detected antibiotic class was tetracycline, comprising ten different genes of which the *tetK* and *tetS* genes were present in five and four samples, respectively. While the *tetK* gene was detected (exclusively) in cheese rind samples containing Staphyloccocal species (*S. aureus, S. saprophyticus* and *S. xylosus*), the *tetS* gene was detected in four samples where streptococcal species were dominant. In particular, *tetS* displays a coverage level similar to *S. thermophilus* genes in C-15, C-18 and C-20, whereas it is 70 times lower than those of MAG-C-13 belonging to *S. salivarius*-like in C-13. Finally, the gene *qnrD1*, detected in the C-R06 and C-R13 samples, was found in contigs with coverage similar to those of *Serratia* and *Providencia* present in both samples, respectively.

**Fig. 5.**
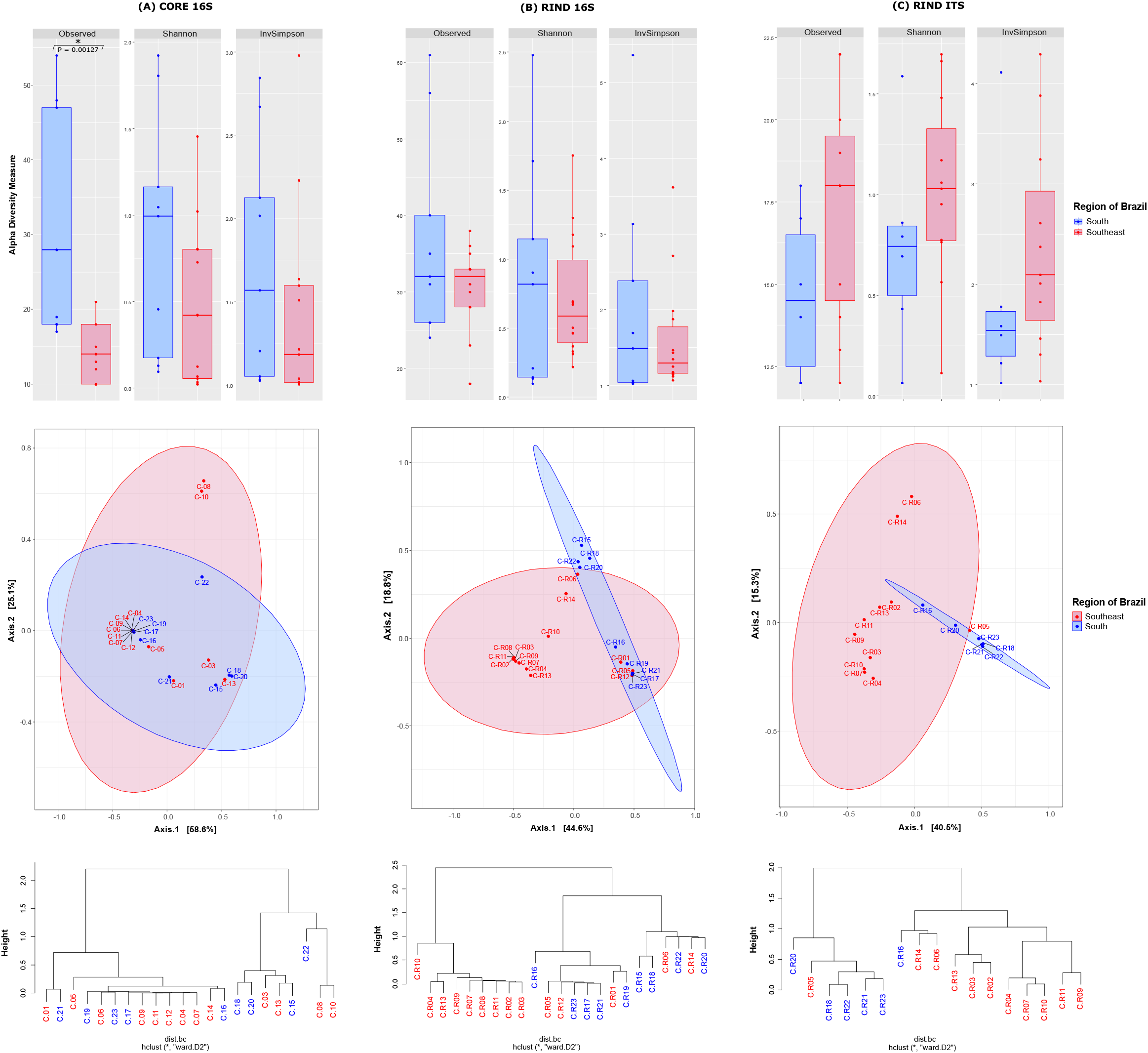
Heatmap showing distribution of the 30 antibiotic resistance genes detected in reads/kb within the 15 metagenomic samples. *MLS: Macrolide, lincosamide and streptogramin.

## 4 Discussion

Artisanal and traditional cheese production has received increased attention as sources of biological resources potentially capable of contributing to technological and health benefits. However, there is an increasing trend, in artisanal production as well, of using starter strains to ensure a rapid and safe lactic fermentation, and, often, the addition of secondary microbiota to more effectively control ripening (Bintsis, 2018; García-Díez & Saraiva, 2021; Vandera, Kakouri, Koukkou, & Samelis, 2019). On the other hand, proper selection of native strains may offer the opportunity to preserve the typicity of the cheese, while offering an excellent level of safety (Gaglio, Todaro, & Settanni, 2021). In order to optimize such a process, cheeses used as original material for strain selection should be carefully selected in order to maximize those with original microbiota and to absolutely avoid those “spoiled” by industrial starter strains, making it difficult to isolate new strains with original properties. In this respect, regions where legislation excludes the use of starter cultures or additives provide the opportunity for the development of autochthonous microorganisms. In this study, we decided to develop a method based on metagenomics to explore the biodiversity of cheese microbiota in order to characterize their originality and potential value as a bio-resource. We studied 23 cheese samples from two regions, South and Southeast Brazil, combining a metagenetic approach to obtain a global view of their microbiota at the genus/family level with metagenomic approaches to access functional and phylogenetic information (**Fig. S5**).

### 4.1 Microbial patrimony of artisanal Brazilian cheeses

As expected, analysis of the cheese cores showed the dominance of LAB such as *L. lactis* and *S. thermophilus*, as already described at the genus level by Kamimura *et al*. (2019) (Kamimura, De Filippis, et al., 2019), except in one sample, which was a production failure, leading to the predominance of an enterobacterium, *Kluyvera cryocrescens*. The analysis of the *L. lactis* MAGs indicated that *L. lactis* present in Brazilian cheese form a particular phylogenomic group within the *lactis* subspecies that differentiate them from the traditional starters (**Fig. S4**). Further analysis such as strain isolation and genome sequencing will be valuable in providing more information about specific genes and should pave the way for technological studies focused on the use of these strains, as biopreservatives, for example, as was shown with the *L. lactis* strain QMF11 isolated from a Brazilian cheese, which has strong anti-listerial activity (Costa et al., 2018).

Interestingly, our study also revealed the presence of two original streptococcal species as the main LABs in several samples. Firstly, *S. infantarius* was found in cheeses from the two regions, and it was even dominant in the core of two SE cheese samples. *S. infantarius* has already been isolated from Brazilian cheeses (Brito et al., 2020; Medeiros et al., 2016) and it is the main LAB isolated from traditional fermented camel and cow milk in East and West Africa, respectively (Jans, Kaindi, et al., 2013). In these countries, the strains isolated from dairy products were shown to contain *lacZ-lacS* genes that share high sequence identity with those of *S. thermophilus*, which provide them with the ability to rapidly assimilate lactose from milk (Jans, Follador, et al., 2013). Further analysis of these sequences showed that East and West African strains independently acquired their *lacZ-lacS* genes from different donor species by horizontal gene transfer (HGT) (Almeida et al., 2014). Our data shows that the Brazilian dairy strains of *S. infantarius* also acquired these genes by HGT from *S. thermophilus* on a large DNA fragment (25,870 nucleotides) flanked by two mobile elements. Features such as the very high identity (99.6%) over the whole length of the fragment and the absence of a recombination event to remove the unnecessary genes indicate that this transfer occurred independently from those described in Africa, and very recently. Secondly, while metagenetic analysis highlighted the presence of Streptococci of the *thermophilus-vestibularis-salivarius* group, which are usually referred to as *S. thermophilus* in dairy products, the metagenomic analysis revealed that cheeses from the Southeast region contain a potentially novel species closely related to *S. salivarius* (**Fig. 4**; **Fig S3A**). The presence in food of these two streptococcal species that probably originated from the human gut, raises the question of their safety in the event of regular consumption. While Streptococci of the *salivarius* group are generally human commensal bacteria that dominate the oral microbiota (Carlsson, Grahnén, Jonsson, & Wikner, 1970; McCarthy, Snyder, & Parker, 1965) and may be used as probiotics (Wescombe, Hale, Heng, & Tagg, 2012), the role of *S. infantarius* in human microbiota is less well established. Nevertheless, its consumption at high levels in African fermented milk indicates that this bacterium is safe for humans (Jans et al., 2017), and a study has shown that it is not associated with colorectal cancer as is its relative in the SBSEC group, *S. gallolyticus* (Jans, Meile, Lacroix, & Stevens, 2015). Overall, these results indicate that cheese-making practices without the use of LAB starters favored the emergence of two novel streptococcal food strains whose properties should be further studied to ensure their safety and to reveal potentially interesting technological properties.

Concerning the cheese rinds, cheeses from the South region, which are not subject to long ripening, mainly contain core bacterial species (**Fig. 3B**), whereas cheeses from the Southeast region are generally dominated by *C. variabile*. Although less prevalent than *C. casei* (Kothe et al., 2021), this halotolerant species is well known for being part of the complex microbiota that develops on the surface of ripened cheeses (Bertuzzi et al., 2018; E. Dugat-Bony et al., 2016) and is also used as a ripening adjunct in several cheese production processes (Delbes, Monnet, & Irlinger, 2015). The *C. variabile-*type strain possesses specific genes associated with metabolic functions involved in the technology and adaptation to cheese habitats (Schröder et al., 2011). As shown by our MAG analysis, *C. variabile* present in SE Brazilian cheeses is closely related to strains isolated from smear ripened-cheeses in Ireland (Schröder et al., 2011) and in France (E. Dugat-Bony et al., 2016). Brazilian cheeses might thus be an interesting source to recover new strains for cheese technology. Moreover, *S. saprophyticus*, a coagulase-negative staphylococcus, was found to be the second most frequent and abundant halotolerant bacteria. This species is frequently detected on the surface of smear-ripened cheese and other fermented foods (Coton et al., 2010; Hammer, Jordan, Jacobs, & Klempt, 2019). While *S. equorum* appears to be the most frequent staphylococcal species in Western cheeses (Kothe et al., 2021), our analysis and an earlier study (Casaes Nunes, Pires de Souza, Pereira, Del Aguila, & Flosi Paschoalin, 2016) indicate that *S. saprophyticus* is more prevalent in Brazilian cheeses. Finally, several less frequent halotolerant species from the genera *Bavaricoccus* and *Brevibacterium* are commonly found in SE cheese, whereas more halophilic species belonging to genera such as *Psychrobacter* and *Halomonas* appear to be scarce compared to Western cheeses (Kothe et al., 2021).

Regarding eukaryotes, overall, we observed that the South samples are dominated by *Debaryomyces hansenii*, while Southeast cheeses are more diverse, notably with the presence of *Diutina catenulata, Kodamae ohmeri, Trichosporon sp*. and *Moniliella sp*. (**Fig. 3C**). Although the presence of these species has been reported worldwide in dairy products (Banjara et al., 2015; E. Dugat-Bony et al., 2016; Irlinger, Layec, Hélinck, & Dugat-Bony, 2015; Wolfe et al., 2014), only *G. candidum* and *D. hansenii* species are known to play a role in cheese ripening (Irlinger et al., 2015; Irlinger & Monnet, 2021; Perkins et al., 2020; Pham, Landaud, Lieben, Bonnarme, & Monnet, 2019). *G. candidum*, in particular, has been found in significant amounts only in one cheese where it is added by the producer, suggesting that this yeast is not naturally of major importance in these artisanal products. Moreover, *Kodamaea ohmeri* and *Diutina catenulata* are poorly studied at the genomic level and no cheese strain genomes of these species are available at this time. The present MAGs thus provide a first insight into their relatedness to environmental yeast strains.

### 4.2 Potential safety concerns

In this study, we determined -through metagenomic reads - that several samples, especially C-13, C-15 and C-18, presented more than 30% of the reads mapped on the *Bos taurus* genome (**Table S3**). Almeida *et al*. (2014) also described similar findings in a blue-veined cheese (Almeida et al., 2014), and the presence at high levels of animal reads in dairy metagenomes could be associated with the presence of milk somatic cells, which are markers of mastitis in dairy cattle, indicating herd health problems (Moradi, Omer, Razavi, Valipour, & Guimarães, 2021; Petzer, Karzis, Donkin, Webb, & Etter, 2017). This may be due to the fact that in artisanal productions, especially in small farms, milk from an animal with mastitis may not be eliminated from the production chain because of the significant financial loss that this may represent.

Regarding microbiological risks, we detected Staphylococci in a number of samples, relatively in accordance with the previous large-scale study of microbial diversity in Brazilian cheeses (Kamimura, De Filippis, et al., 2019). Since this study was made at the genus level, it left open the possibility of a high prevalence in Brazilian cheeses of *S. aureus*, a common pathogen in dairy industries worldwide (Cretenet, Even, & Le Loir, 2011), including in Brazil (Dittmann et al., 2017). We were able to show with a high degree of confidence that most species belong to coagulase-negative species such as *S. saprophyticus*, whereas a significant amount of *S. aureus* was detected in only one sample. However, in this sample, we identified the presence of the heat-stable enterotoxin A (*sea*), which is associated with illness, accounting for 77.8% of all staphylococcal foodborne disease outbreaks (Argudín, Mendoza, & Rodicio, 2010; Balaban & Rasooly, 2000; Kadariya, Smith, & Thapaliya, 2014), indicating a potential health risk to consumers.

Furthermore, we detected the presence of several transmissible antibiotic resistance genes that are considered to be a growing threat to public health, both in hospital and food industry environments (Y. Li et al., 2020; Yadav & Kapley, 2021). These notably include *tetS* and *tetK* genes, which are probably associated with the presence of streptococcal and staphylococcal species, respectively. Tetracycline resistance has been largely reported in fermented foods, for example, in cheese ripening (Ana Belén Flórez, Delgado, & Mayo, 2005; Ana B. Flórez, Vázquez, & Mayo, 2017; X. Li, Li, Alvarez, Harper, & Wang, 2011) and, more specifically, the occurrence of the *tetS* gene has been associated with *S. thermophilus* in cheeses (Ge et al., 2007; Rizzotti, La Gioia, Dellaglio, & Torriani, 2009), whereas that of *tetK* has been associated with *S. aureus* in dairy products (Jamali, Paydar, Radmehr, Ismail, & Dadrasnia, 2015) and different species of coagulase-negative staphylococci in salami samples (Rebecchi, Pisacane, Callegari, Puglisi, & Morelli, 2015). The presence of antibiotic resistance genes at high levels is often associated with the use of milk from animals treated with antibiotics (Tóth et al., 2020).

## 5 Conclusion

The data presented here show that the artisanal cheeses produced in the South and Southeast regions of Brazil display an original microbiota, only marginally contaminated by industrial starters. Interestingly, it contains original lineages of bacterial strains belonging to known species of technological interest, such as *L. lactis* and *C. variabile*, as well as yeasts such as *D. hansenii*. It also contains less ordinary food species such as the bacteria *B. seileri, B. alimentarium, S. saprophyticus*, and the yeast *D. catenulata*. Moreover, two new food streptococci, *S. infantarius* and *S. salivarius-like* were found to be the main LAB in several production schemes, confirming the high originality of the Brazilian artisanal cheese microbiota and the value of this small-scale production scheme, which collectively represent a reservoir of biodiversity. Additional studies should be performed in order to improve our understanding of these microbiota and their impact on the sensory aspects of Brazilian cheeses. The further characterization of representative strains will also open the possibility to develop inoculants that maintain the cultural and historical identities of these cheeses and to establish standards of quality for their production. Finally, our study identifies several areas of concern, such as the presence of somatic cell DNA and high levels of antibiotic resistance genes in several microbiota, inferring that the milk used was from diseased herds. Overall, the data from this study highlight the potential value of the traditional and artisanal cheese production network in Brazil, and provide a metagenomic-based scheme to help manage this resource safely.

## Authors’ contributions

CIK and PR conceived the study and its experimental design. CIK performed microbiological, genomic, metagenomic and functional analyses. NM and PR contributed to genomic and functional analysis. CIK provided data visualization. CIK and PR wrote the manuscript. PR supervised the project.

## Acknowledgements

The authors are grateful to Mathieu Almeida and Stéphane Chaillou for sharing their knowledge with us about metagenomic data analysis.

CIK’s grant was supported by the “Conselho Nacional de Desenvolvimento Científico e Tecnológico (CNPq)” – Brazil [grant number 202444/2017-1].

## Supplementary Material

**Supplementary Data 1**. The production process and characteristics of Araxá, Canastra, Serro, Colonial and Serrano.

**Fig. S1**. Rarefaction curves depicting the depth of 16S (core and rind) and ITS (rind) sequencing, as well as species richness for the data obtained from Brazilian cheeses. The x-axis represents the sequencing depth (reads) and the y-axis represents the estimated OTU richness detected at species level.

**Fig S2**. Phylogeny of species detected by *ychF*, highlighting the reconstituted MAGs.

**Fig. S3**. Potential new species and genera found by autoMLST analyses.

**Fig. S4**. Phylogenomic tree for *Lactococcus lactis* highlighting the different groups of *L. lactis*.

**Fig. S5**. Overview of approaches used in this study.

**Table S1**. Metadata describing the 23 cheese samples analyzed.

**Table S2**. Raw reads detected of bacterial and fungal operational taxonomic units (OTUs) in the samples.

**Table S3**. Percentage (%) of *Bos Taurus* reads aligned in each cheese metagenome.

**Table S4**. The relative abundance of microbial composition of 15 cheese metagenomes from Brazil using MetaPhlAn taxonomic assignment and *ychF* marker gene methods.

**Table S5**. Quality of prokaryotic and eukaryotic MAGs and their ANIs with the closest reference genome found in the NCBI database.

**Table S6**. Coverage (reads/kb) of antibiotic resistance and genes of technological interest. **Table S7**. Detection of 12 pathogens commonly found in dairy environments by read mapping expressed in percentage of total read mapped on a set of reference genomes.

