## Supplementary figures and images for "Revealing the microbial heritage of traditional Brazilian cheeses through metagenomics"

### Figure S1

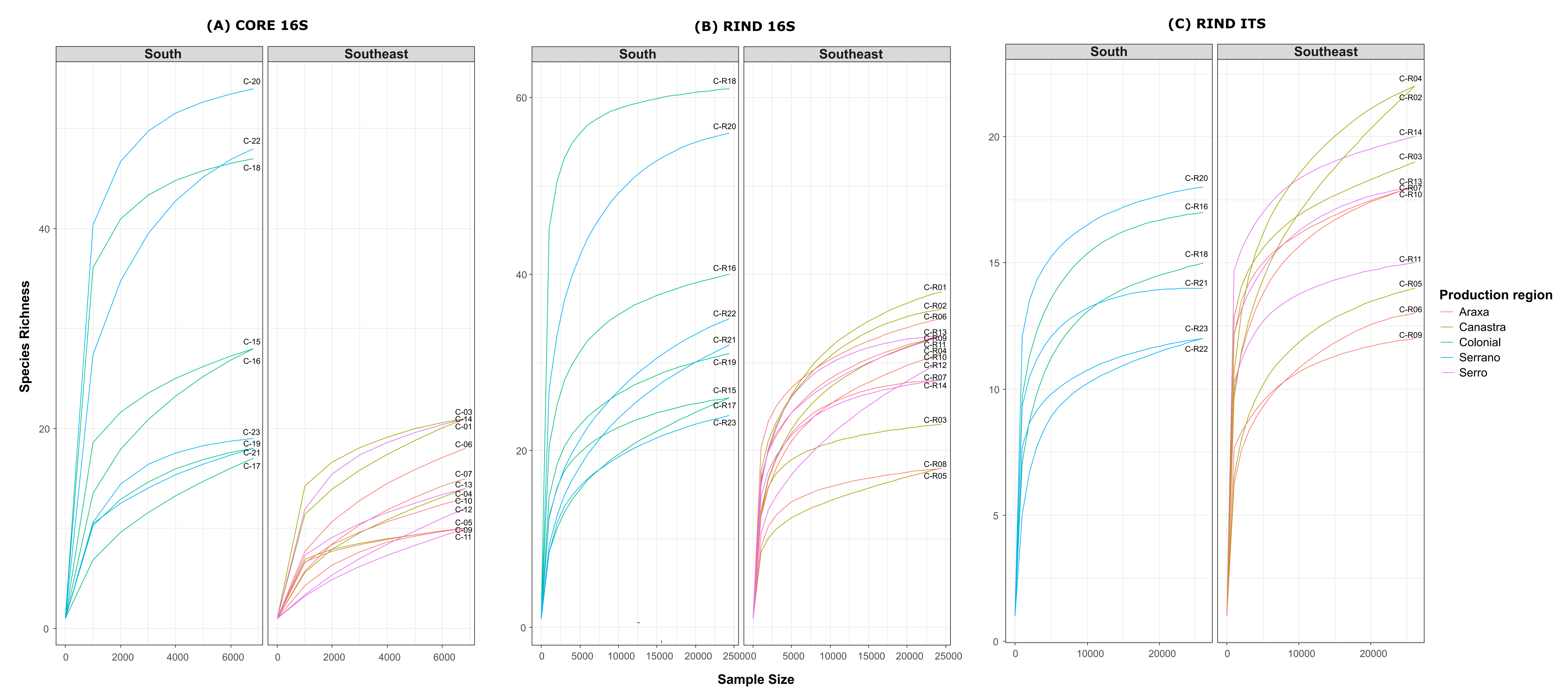

### Figure S2

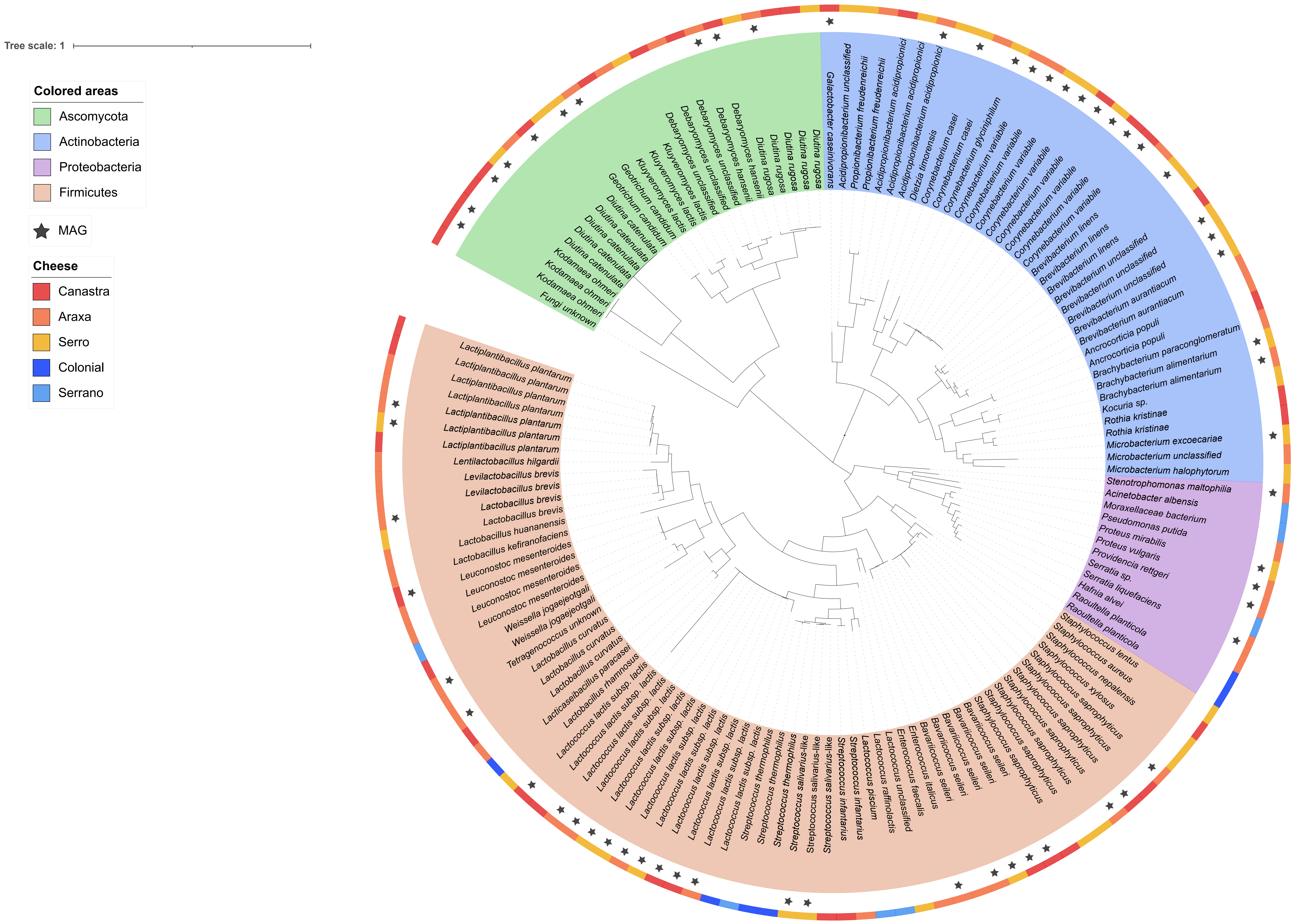

### Figure S3

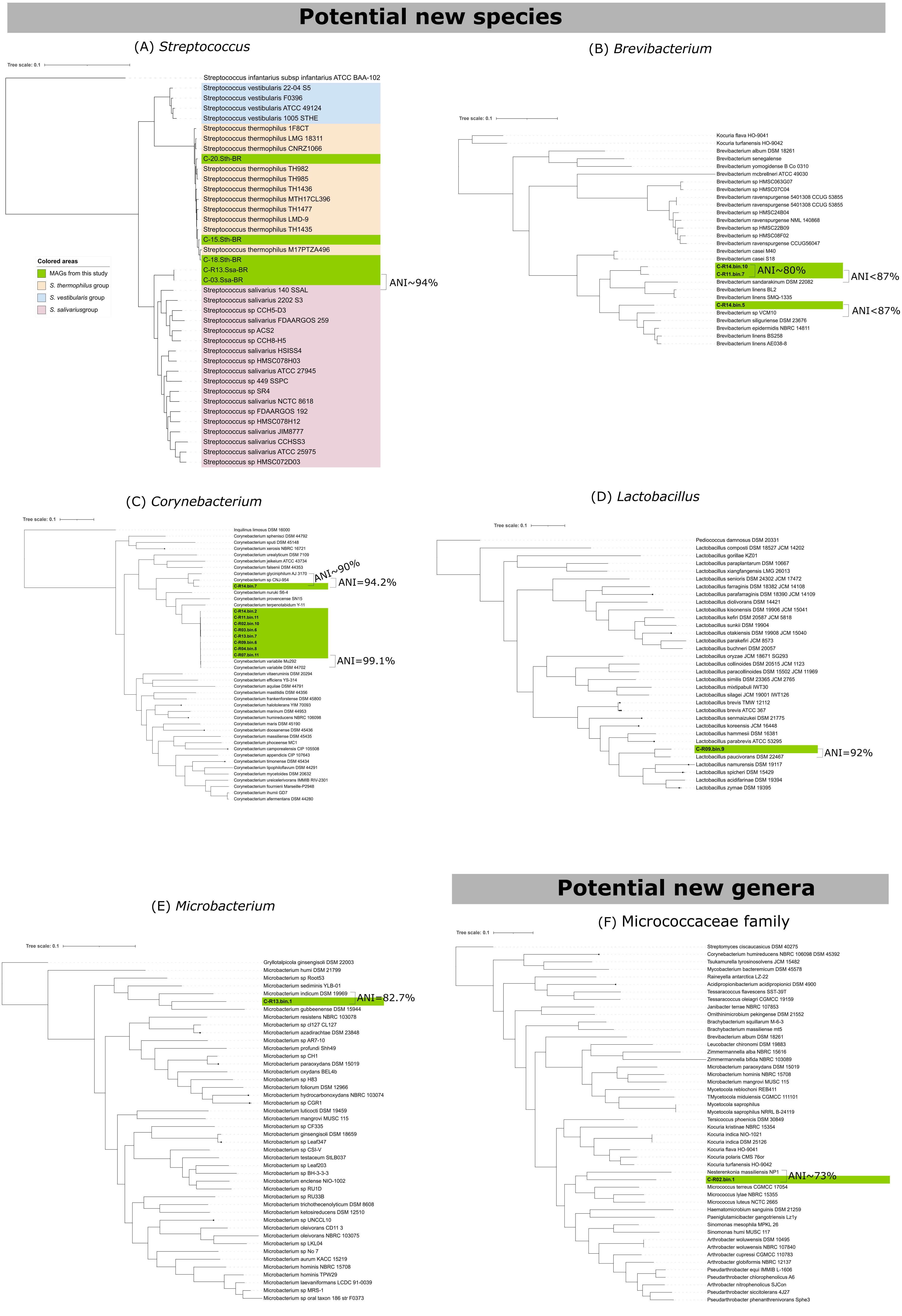

### Figure S4

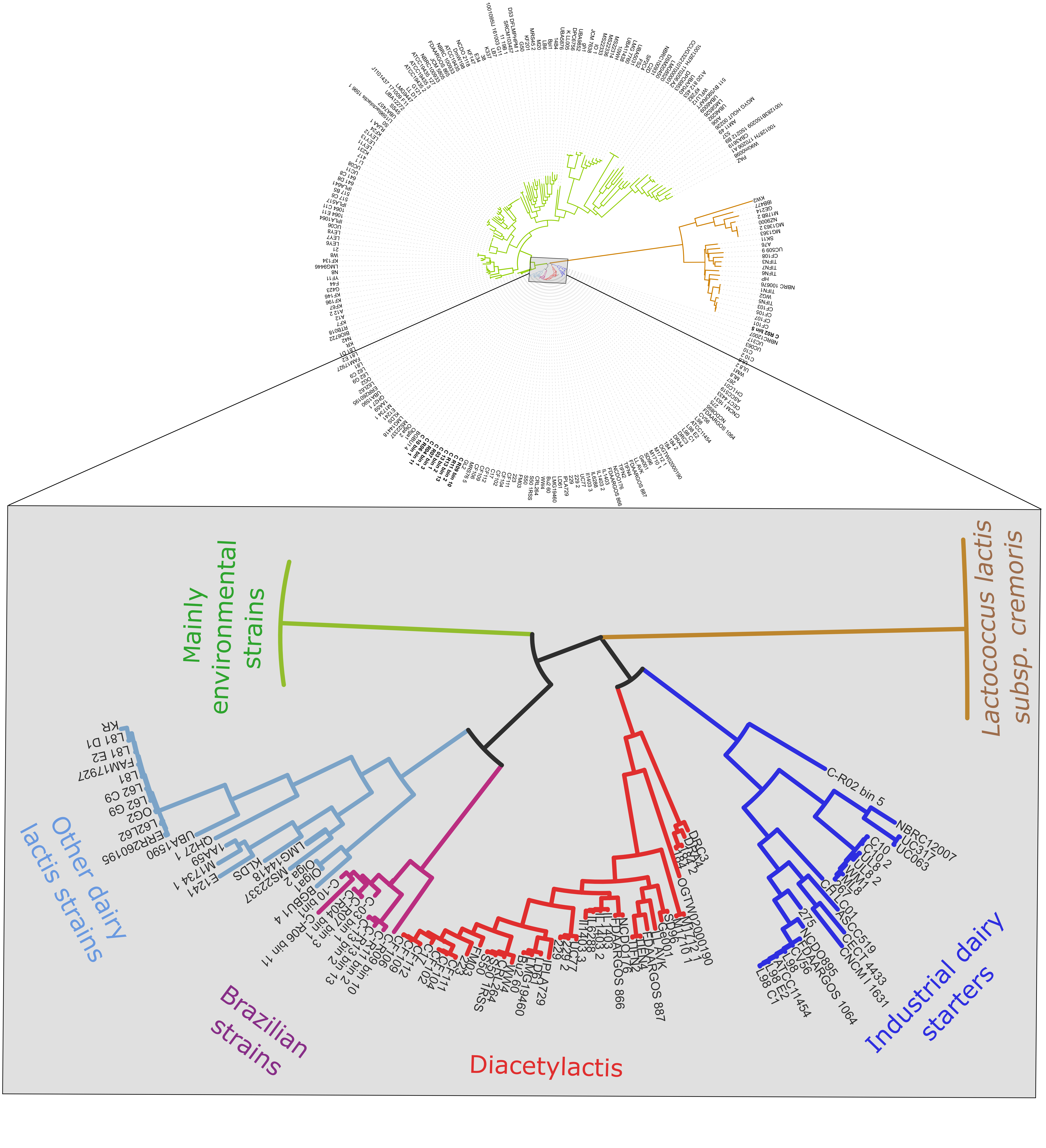

### Figure S5

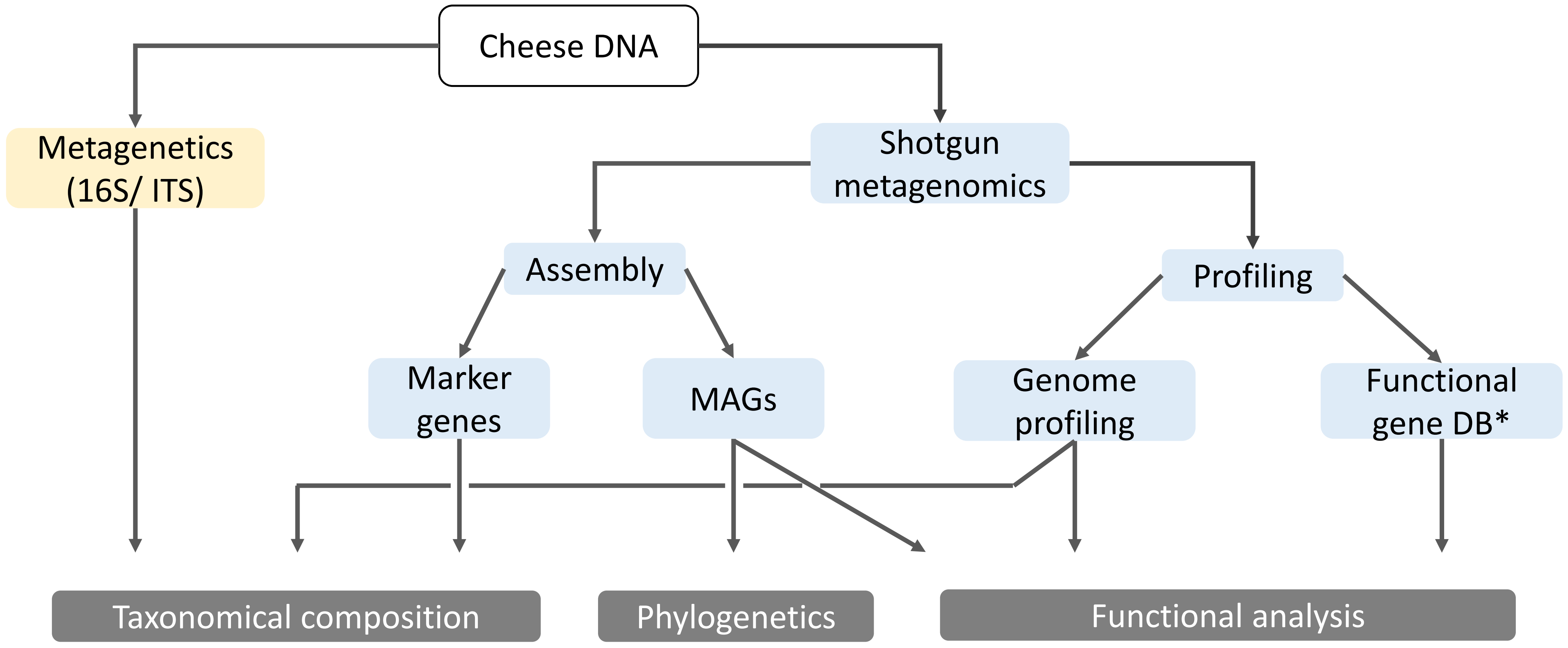
