## Supplementary Data 1 for "Revealing the microbial heritage of traditional Brazilian cheeses through metagenomics"

Colonial and Serrano cheeses are manufactured in the South of Brazil, in the states of Rio Grande do Sul (RS) and Santa Catarina (SC). Their processing includes the basic steps for cheese-making: milk filtration, addition of salt to milk or to the curd, addition of rennet, coagulation, cutting of the curd, draining, molding, pressing and ripening (Kamimura et al., 2019). Although there are no standardization or regulation techniques for the production of Colonial cheese in Brazil (de Medeiros Carvalho, de Fariña, Strongin, Ferreira, & Lindner, 2019), it is usually made from the milk of Jersey or Dutch dairy cows and has a heating step (35-40°C) after the curd has been cut (Fava, Hernandes, Pinto, & Schmidt, 2012; Kamimura et al., 2019). It is a cheese with some eyes, a yellowish color, marketed in pieces ranging from 0.7 to 5 kg, and with a ripening period that varies between 30 and 75 days (Fava et al., 2012; Kamimura et al., 2019). Meanwhile, in 2016, a regulation on the production and commercialization of Serrano cheese was approved in RS (RS, 2016). This cheese is made using raw milk from beef cattle breeds, which produce reduced amounts of milk with high fat concentration (Delamare, C P Andrade, Mandelli, Almeida, & Echeverrigaray, 2012; MAPA, 1996). The use of commercial lactic cultures, preservatives and additives is forbidden in the manufacture of Serrano. There is no heating stage and manual pressing is carried out before molding (Amarante, 2015; Kamimura et al., 2019). Serrano cheese is generally consumed after a short ripening period (15-30 days) and is available in round or rectangular shapes (around 2-3kg) (Borelli, Lacerda, Penido, & Rosa, 2016).

Araxá, Canastra and Serro cheeses are manufactured in micro-regions in Minas Gerais (MG) state, the largest producer of cheeses in Brazil and the region with the oldest regularization for the production and commercialization of artisanal cheeses (MG, 2002). The main difference in the processing of these cheeses is the use of “pingo”, a natural starter obtained from cheese whey, produced after molding, which is added before the coagulation step (Perin, Sardaro, Nero, Neviani, & Gatti, 2017). Although the stages of the cheese-making process are similar among the micro-regions, the climatic factors (altitude, temperature and relative humidity of the air) could favor the development of a unique identity, despite the absence of documented differences (Borelli et al., 2016; Kamimura et al., 2019). The main difference in the production process between these three artisanal cheeses from MG is that those made in the Serro micro-regions are pressed manually (using the hands), while the cheeses made in the Canastra and Araxá micro-regions are pressed with the help of a cheesecloth (Kamimura et al., 2019). Additionally, the ripening period is defined as a minimum of 14 days for Araxá, 17 days for Serro, and 22 days for Canastra (IMA, 2017).
