## Supplementary material for "Revealing the microbial heritage of traditional Brazilian cheeses through metagenomics": Table S1

| **Sample** | **Cheese** | **Region of Brazil** | **Milk** | **Collected** |
| --- | --- | --- | --- | --- |
| C-01 | Canastra | Southeast | Raw | Artisan market |
| C-02 | Canastra | Southeast | Raw | Artisan market |
| C-03 | Canastra | Southeast | Raw | Artisan market |
| C-04 | Canastra | Southeast | Raw | Artisan market |
| C-05 | Canastra | Southeast | Raw | Artisan market |
| C-06 | Araxá* | Southeast | Raw | Artisan market |
| C-07 | Araxá | Southeast | Raw | Artisan market |
| C-08 | Araxá | Southeast | Raw | Producer |
| C-09 | Araxá | Southeast | Raw | Artisan market |
| C-10 | Araxá | Southeast | Raw | Artisan market |
| C-11 | Serro | Southeast | Raw | Artisan market |
| C-12 | Serro | Southeast | Raw | Artisan market |
| C-13 | Serro | Southeast | Raw | Artisan market |
| C-14 | Serro | Southeast | Raw | Producer |
| C-15 | Colonial | South | Raw | Artisan market |
| C-16 | Colonial | South | Raw | Producer |
| C-17 | Colonial | South | Raw | Artisan market |
| C-18 | Colonial | South | Raw | Artisan market |
| C-19 | Colonial | South | Pasteurized | Public market |
| C-20 | Serrano | South | Raw | Producer |
| C-21 | Serrano | South | Raw | Fair |
| C-22 | Serrano | South | Raw | Fair |
| C-23 | Serrano | South | Raw | Public market |

* Addition of *Geothricum candidum* indicated on the label
